# Constant conflict between *Gypsy* LTR retrotransposons and CHH methylation within a stress-adapted mangrove genome

**DOI:** 10.1101/263830

**Authors:** Yushuai Wang, Weiqi Liang, Tian Tang

## Abstract

**Summary:**

- Evolutionary dynamics of the conflict between transposable elements (TEs) and their host genome remain elusive. This conflict would be intense in stress-adapted plants as stress can often reactivate TEs. Mangroves reduce TE load convergently in their adaptation to intertidal environments and thus provide a unique opportunity to address the host-TE conflict and its interaction with stress adaptation.
- Using the mangrove *Rhizophora apiculata* as a model, we investigated methylation and short interfering RNA (siRNA) targeting patterns in relation to the abundance and age of long terminal repeat (LTR) retrotransposons. We also examined LTR retrotransposons’ distance to genes, impact on neighboring gene expression, and population frequencies.
- We found differential accumulation among classes of LTR retrotransposons despite high overall methylation levels. This can be attributed to 24-nt siRNA-mediated CHH methylation preferentially targeting *Gypsy* elements, particularly in their LTR regions. Old *Gypsy* elements possess unusually abundant siRNAs which show cross-mapping to young copies. *Gypsy* elements appear to be closer to genes and under stronger purifying selection than other classes.
- Our results suggest a continuous host-TE battle masked by the TE load reduction in *R. apiculata*. This conflict may enable mangroves like *R. apiculata* to maintain genetic diversity and thus evolutionary potential during stress adaptation.

## Introduction

Plants suppress transposable element (TE) activity through RNA interference (RNAi) and DNA methylation. Post-transcriptional silencing of transposons is triggered by 21-22 nucleotide (nt) short interfering RNAs (siRNAs) which guide cleavage of TE mRNAs. Transcriptional silencing of TEs via methylation occurs in three sequence contexts: CG, CHG and CHH (where H stands for A, T or C) (Cokus *et al.*, 2008; Lister *et al.*, 2008). While CG and CHG methylation can be maintained and inherited by daughter DNA strands (Kankel *et al.*, 2003), CHH methylation relies on *de novo* targeting of RNA-directed DNA methylation (RdDM) (Zemach *et al.*, 2013). The targeting of TEs for *de novo* cytosine methylation is usually mediated by 24-nt siRNAs (Wassenegger *et al.*, 1994; Pontier *et al.*, 2012). Transposon methylation can negatively affect expression of neighboring genes and thus potentially contributes to the deleterious effects of these elements, resulting in purifying selection against insertion events (Hollister & Gaut, 2009).

Evolutionary dynamics of the conflict between TEs and their host genomes remain elusive. Such conflict is expected to be intense in plants growing in extreme environments. On one hand, TEs are often activated under stressful environments, providing extensive genetic diversity for selection to act on (Kidwell & Lisch, 1997; Chenais *et al.*, 2012; Casacuberta & Gonzalez, 2013). On the other hand, TE load might become a heavy burden for hosts that struggle to survive in hostile environments. This dilemma is part of a fundamental debate that seeks to establish whether and how adaptively optimal mutation rates are achieved by organisms. Comparative analyses between *Arabidopsis* species and natural populations of wild barley inhabiting contrasting habitats often find taxa that thrive in harsh environments carry heavy TE burdens and large genome sizes (Hollister *et al.*, 2011; Wu *et al.*, 2012; Kalendar *et al.*, 2000). It is thus surprising that the recent findings in mangroves paint a different picture.

Mangroves independently invaded tropical and subtropical intertidal swamps that provide extremely stressful environments for tree species (Tomlinson, 1994). This change in lifestyle, together with availability of closely-related non-mangrove species to serve as controls, provides a good opportunity to study evolutionary consequences of interactions between TE load and host stress. A recent study found massive and convergent TE load reduction in mangroves in comparison to non-mangrove species (Lyu *et al.*, 2018). This reduction was attributed to the paucity of young long terminal repeat (LTR) retrotransposons as a consequence of fewer births rather than excess death (Lyu *et al.*, 2018). In the present study, we investigated methylation and siRNA targeting patterns in the mangrove *Rhizophora apiculata* and related these patterns to the abundance and age of LTR retrotransposons. Our results indicate preferential suppression of active and thus most dangerous TE elements via 24-nt siRNA-mediated CHH methylation in *R. apiculata*. We also show evidence that reactivation of old TEs can trigger rapid silencing of young copies from the same family. The constant host-TE conflict may reflect a balance between alleviating the TE burden and maintaining genetic diversity as mangroves adapt to stressful environments.

## Materials and Methods

### Annotation of *R. apiculata* LTR retrotransposons

The PacBio and Illumina hybrid assembly of the *R. apiculata* genome was retrieved from the European Nucleotide Archive (ENA) [accession number PRJEB8423, (Xu *et al.*, 2017)]. LTR_Finder (Xu & Wang, 2007) and LTR_harvest (Ellinghaus *et al.*, 2008) were used to *de novo* search for LTR retrotransposons in the *R. apiculata* genome with default parameters. Predictions by these two work flows were merged and filtered by genomic coordinates requiring at least 1 kb distance between two adjacent candidates. We then annotated the internal sequences of candidate LTR retrotransposons using BLASTX against profile Hidden Markov Model gene models from the Gypsy Database 2.0 (Llorens *et al.*, 2011) with the e-value cutoff 1e-8. Candidate LTR retrotransposons that contain typical Gag-Pol protein sequences were retained as intact elements. All the intact LTR retrotransposons were clustered using SiLiX (Miele *et al.*, 2011) and considered as a family if they were ≥80% identical over ≥80% of their aligned sequence of at least 80 bp in length (Wicker *et al.*, 2007). All intact LTR retrotransposons were classified into *Copia*, *Gypsy,* and unclassified groups according to the order of identified protein domains (Wicker *et al.*, 2007).

Once we finalized our LTR retrotransposon family assignments, we conducted a second round of annotation using intact elements as queries to search against the *R. apiculata* genome. LTRs and internal sequences from each family were subject to BLASTN separately. The minimum aligned length and e-value cutoff were ≥70% and 1e-8 for LTRs, and ≥50% and 1e-8 for internal regions. Combining the genomic coordinates, we obtained a full list of intact elements (Table S1), truncated elements and solo-LTRs for each family.

### Age estimation and transition:transversion ratios

LTR retrotransposon age was estimated as previously described (SanMiguel *et al.*, 1998). The LTR pairs from each TE insertion were aligned using MUSCLE (v3.8.31) with default parameters (Edgar, 2004). Kimura’s two-parameter model (Kimura, 1980) was used to estimate sequence divergence *k* between pairs of LTRs. The age of LTR retrotransposons was estimated using the formula T=*k*/2*r*, where *r* is the substitution rate of 1.3×10^−8^ substitutions/site/year (Ma & Bennetzen, 2004).

The transition (Ts) and transversion (Tv) mutation counts between LTR pairs were estimated using Kimura’s two-parameter model (Kimura, 1980). The ratio of these numbers is the transition: transversion (Ts:Tv) ratio. For LTR retrotransposons from a specific group or species, the Ts and Tv numbers were summed up separately to obtain an average Ts:Tv ratio. Genomic sequences of *Arabidopsis thaliana* and *Populus trichocarpa* were retrieved from Phytozome v12.1 (https://phytozome.jgi.doe.gov). We used TE annotations as described by TAIR10 (https://www.arabidopsis.org) and RepPop (Zhou & Xu, 2009).

### Bisulfite sequencing and methylome analyses

Young leaves of *R. apiculata* were collected from Qinlan Harbour, Hainan, China for whole-genome bisulfite sequencing. Genomic DNA was extracted using the modified CTAB protocol (Doyle & Doyle, 1987). Bisulfite treatment of genomic DNA was performed using the Zymo EZ DNA Methylation Lightning Kit (Zymo Research). The bisulfite-treated DNA was purified and used to prepare sequencing libraries using the EpiGnome™ Kit (Epicentre). The libraries were sequenced on an Illumina HiSeq 2500 using 100 bp paired-end reads. All procedures followed the manufacturers’ instructions.

Bisulfite sequencing reads were trimmed to get rid of adapters and low-quality bases using trimmomatic v0.32 (Bolger *et al.*, 2014), and then mapped to the *R. apiculata* genome using Bismark ver. 0.16.3 (Krueger & Andrews, 2011) with default parameters. Only uniquely mapping reads were retained. The conversion rate (rate at which unmethylated cytosines were converted to uracil by bisulfite) was calculated by using reads mapping to the lambda genome. Cytosines were called as methylated (FDR < 0.05) using a binomial test using the conversion rate as the expected probability followed by multiple test correction using the Benjamini–Hochberg false discovery rate (FDR). Each LTR retrotransposon and its two kilobase upstream and downstream regions was divided into 20 equally-sized windows. Methylation levels of specific windows were calculated as the proportion of Cs among Cs and Ts (#C/(#C+#T) by methylation context (CG, CHG and CHH) using custom Perl scripts. Only sites covered by more than three reads were used for the calculation (Schultz *et al.*, 2012). Correlation between methylation level and TE age was estimated in R v3.1.3.

### Small RNA sequencing and data analysis

Total RNA was extracted from young *R. apiculata* leaves using a modified CTAB method (Yang *et al.*, 2008). Small RNA library construction and sequencing was conducted as previously described (Wen *et al.*, 2016). After adaptor trimming and quality control, clean reads ranging from 18 to 30 nt were aligned against Rfam (Rfam v. 13.0) and miRbase v. 21 (Kozomara & Griffiths-Jones, 2014) using Bowtie (Langmead *et al.*, 2009) to remove structural non-coding RNAs, such as rRNA, tRNA, snRNA, snoRNA, and known miRNAs. The remaining reads were considered to be putative siRNAs and aligned to the *R. apiculata* genome using Bowtie (Langmead *et al.*, 2009). No mismatches were allowed in any of the steps. siRNA expression levels at LTR retrotransposons were calculated as read counts per base pair per element. If a siRNA had multiple hits to different positions in the genome, we assigned the total read count divided by the number of hits to each location and added these data points to the uniquely mapping siRNAs. Correlation between siRNA expression and TE age was estimated in R v3.1.3.

### Sequence divergence between and within groups of LTR retrotransposons

DNA sequences of LTRs and the internal region of each retrotransposon family were extracted and aligned separately using MUSCLE v3.8.31 (Edgar, 2004). Coding sequences were aligned with the guide of amino acid sequences and adjusted manually. The synonymous substitution rate (Ks) in pairwise comparisons was calculated using baseml from the PAML package (Yang, 2007) using the K80 model. Ks of the 5’ and 3’ LTR sequences were calculated separately and the average Ks was used to represent the divergence between LTRs.

### Transcriptome sequencing and expression of TEs and TE-neighboring genes

Total RNA from *R. apiculata* leaves was used for RNA-seq using pair-end 150-bp reads. Two biological replicates were sequenced. After adapter-trimming and quality control using FastQC v0.10.1 and trimmomatic v0.32 (Bolger *et al.*, 2014), clean reads were mapped to intact LTR retrotransposons using Bowtie (Langmead et al., 2009) allowing no more than two mismatches. Expression level of each LTR retrotransposon copy was calculated as Reads Per Kilobase of transcript and averaged between the two biological replicates. To measure gene expression, clean reads were mapped to the repeat-masked *R. apiculata* genome using Bowtie (Langmead *et al.*, 2009) with no more than two mismatches, and analyzed using HTSeq ver. 0.6.1 (Anders *et al.*, 2015) with default parameters. Expression level of each gene was calculated as Reads Per Kilobase per Million mapped reads (RPKM) and averaged between the two biological replicates. TEs or genes with no reads mapping in both replicates were discarded in further expression analyses. We measured the distance from a TE to its nearest neighboring gene, including both upstream and downstream genes, as described previously (Hollister & Gaut, 2009).

### Whole-genome population sequencing and estimation of TE insertion frequency

We sampled an *R. apiculata* population in Qinlan Harbour, Hainan, China. Equal amounts of silica-gel-dried leaves from 10 individuals were pooled for DNA extraction and library construction. Whole genome sequencing was conducted on an Illumina HiSeq 2000 using 125-bp paired-end reads. Overall, the pool-seq data had ~300-fold coverage. We used T-lex2 (Fiston-Lavier *et al.*, 2015) to estimate population frequencies of LTR retrotransposon insertions that were annotated in the reference genome. TE frequency was estimated as the ratio of the number of sequence reads that support the presence of the insertion to the total number of reads covering the genomic site.

Using polymerase chain reaction (PCR), we evaluated the presence/absence status of 23 randomly selected TE insertions that were polymorphic in the sequencing pool in additional 32 individuals from the same population. The multiplex PCR for each insertion was designed as previously described (Gonzalez *et al.*, 2008). Two sets of primers were used, one located in the 5’ (F1) and 3’ flanking regions (R) of the insertion to assay the presence of the insertion, the other located within the TE (F2) and 3’ flanking region (R) to assay the absence of the insertion (Fig. S6a). All PCR primers were designed using Primer 3 (Untergasser *et al.*, 2012) and are listed in Table S3. PCR reactions were conducted using 2×Taq PCR StarMix with Loading Dye (GenStar) following the manufacturer’s instructions. Population frequency of each insertion was estimated using a maximum likelihood procedure (Gonzalez *et al.*, 2008). Data generated in this study were deposited in the Short Read Archive Database under the accession number PRJNA432798.

## Results

### Elevated transition/transversion ratios in mangrove LTRs

The recently completed *R. apiculata* genome is ~274 Mb long (Xu *et al.*, 2017). The hybrid PacBio and Illumina assembly contains 142 scaffolds with N50 of 5.4 Mb (Xu *et al.*, 2017). We annotated intact LTR retrotransposons in this sequence and clustered them into families (see Materials and Method). We found 1,446 intact LTR retrotransposons, comprising 30 families of 1,068 *Copia* elements, 10 families of 278 *Gypsy* elements, and 12 families of 100 unclassified elements (Fig. 1a and Table S1). The copy number of *Copia* elements far exceeded the other two groups. The top three most abundant *Copia* families accounted for more than 49% of the total number of intact LTR retrotransposon copies (Fig. 1a). It appears that LTR retrotransposon groups in the *R. apiculata* genome accumulate at different rates.

**Fig. 1.**
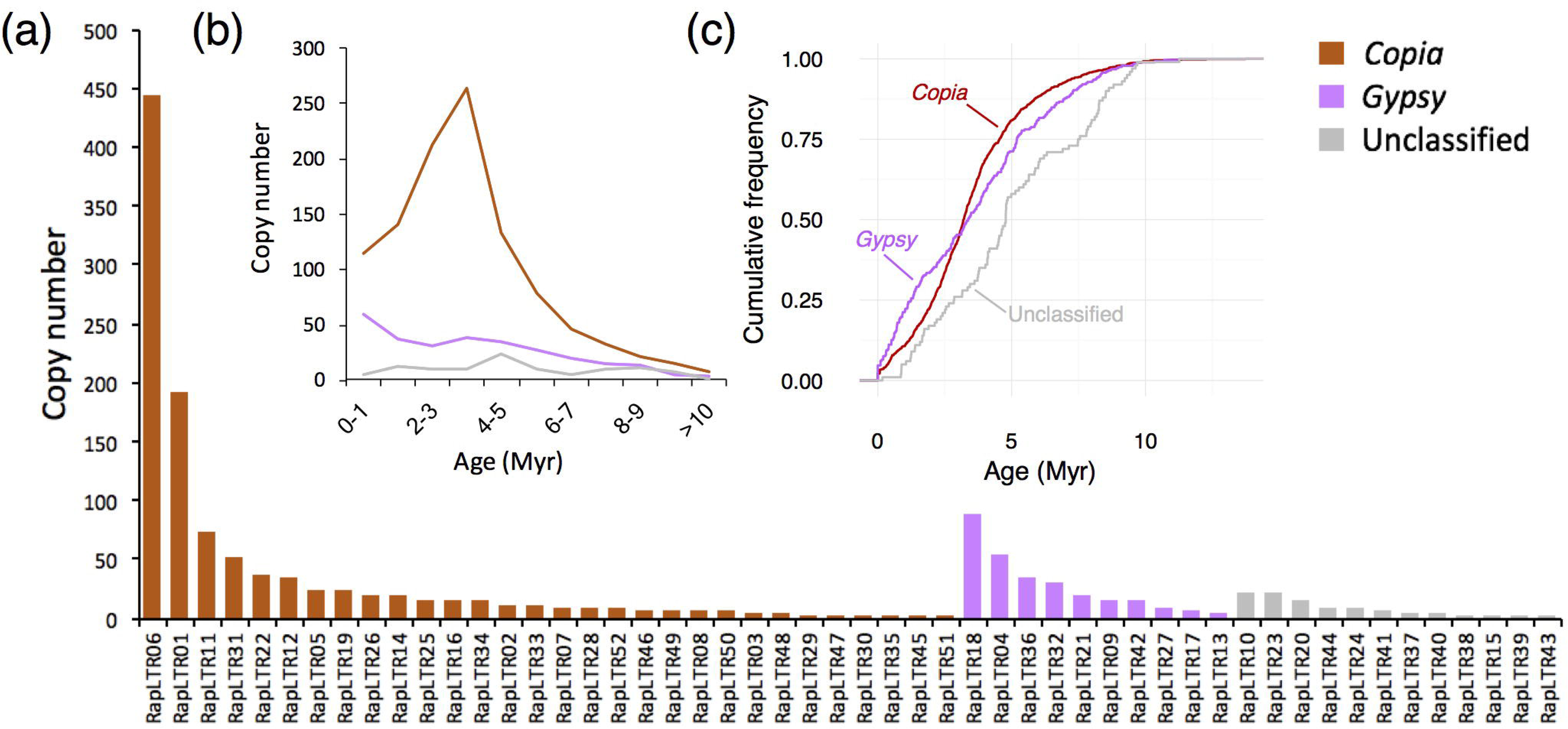
Copy number and age distribution of *R. apiculata* LTR retrotransposon families. (a) Distribution of the number of elements across LTR retrotransposon families. (b) Line graph showing age distributions of three groups of LTR retrotransposons. (c) Cumulative plot of age distributions of three groups of LTR retrotransposons. The *Copia*, *Gypsy*, and unclassified groups are indicated in brown, pink, and grey, respectively.

We asked whether DNA methylation, a major form of epigenetic regulation, may affect TE abundance in *R. apiculata*. Methylated cytosine residues spontaneously deaminate to form thymine residues, leading to an elevated ratio of transitions to transversions (Ts:Tv) between the two LTRs residing at the ends of individual retrotransposons (Vitte & Bennetzen, 2006; Baucom *et al.*, 2009). If strong methylation accounts for the rarity of young LTR retrotransposons in mangroves, we would expect to see higher Ts:Tv ratios in mangrove TEs compared to those in closely-related non-mangrove species. Indeed, the global Ts:Tv ratio, at 5.16, in the LTRs of retrotransposons is three to six fold higher in *R. apiculata* than in *Populus trichocarpa* (0.85) or *Arabidopsis thaliana* (1.6) (Table 1). TEs account for 7.83% of the *R. apiculata* genome (Lyu et al. 2017) whereas the fraction is approximately 40% in *P. populus* (Zhou & Xu 2009) and ~10% in *Arabidopsis* (The *Arabidopsis* Genome Initiative, 2000). In contrast, Ts:Tv ratios are comparable between the *Copia* and *Gypsy* elements within *R. apiculata* (5.47 vs. 5.06), even though the abundance of retrotransposon groups is markedly different. Both *Copia* and *Gypsy* elements exhibit a higher Ts:Tv ratio than unclassified elements (3.20) (Table 1). When Ts:Tv ratios were calculated for individual TE families, we found a positive correlation between these ratios and TE copy numbers (Pearson’s correlation, *r* = 0.36, P < 0.05). Although *Copia* families on average have a higher Ts:Tv ratio than *Gypsy* families (4.26 vs. 4.75), the difference is not statistically significant (*t*-test, *P*>0.05).

**Table 1.**
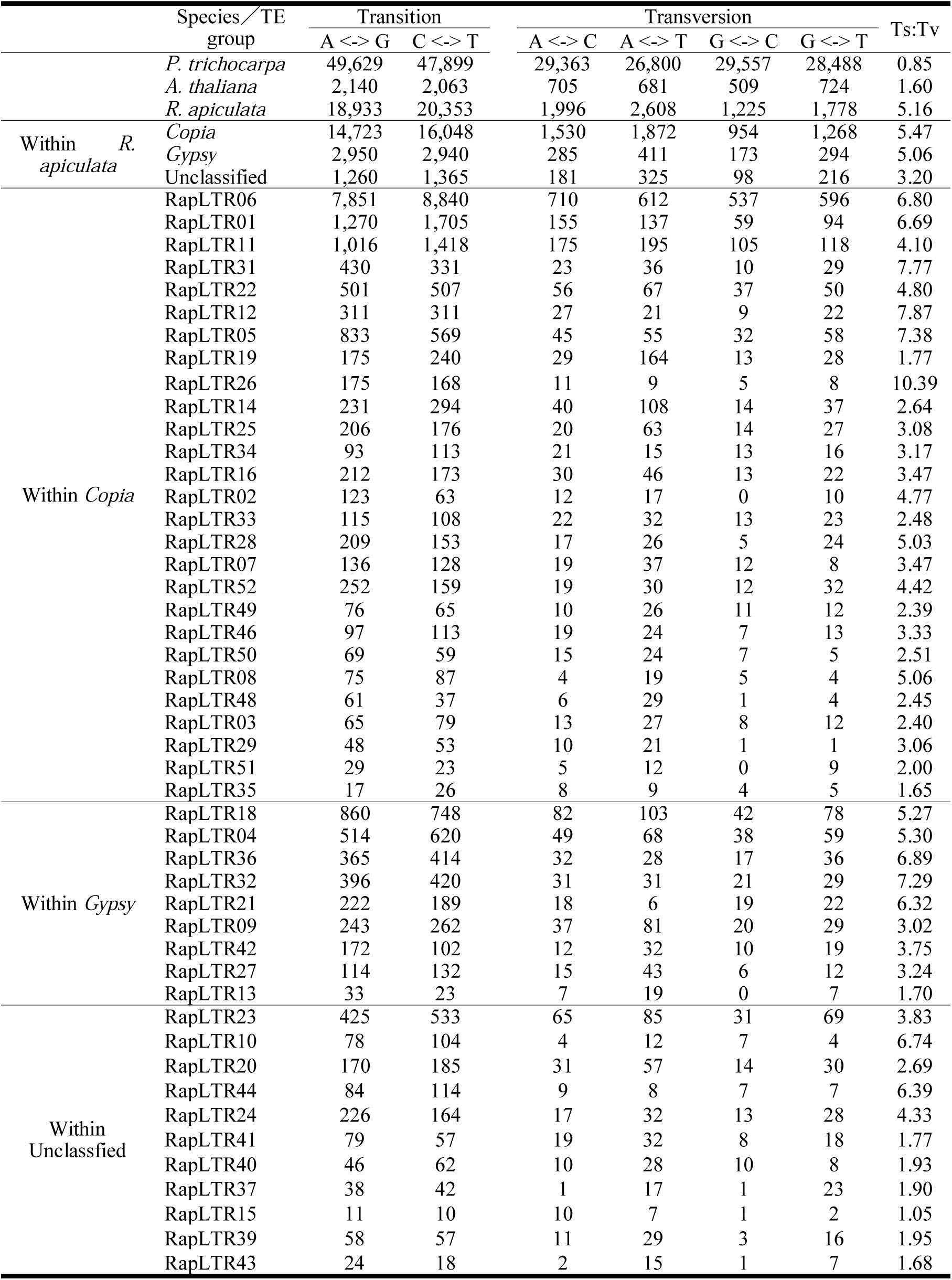
Transition to transversion ratios (Ts:Tv) in LTR retrotransposons from specific species, groups within the *R. apiculata* genome, or families within each LTR retrotransposon group of *R. apiculata*.

### Differential TE accumulation correlates with CHH methylation that is predominantly in LTRs

DNA methylation levels change as a function of TE age. We suspected that age-related methylation may account for the differential TE accumulation within the *R. apiculata* genome. To investigate DNA methylation as a function of TE age, we estimated ages of intact elements by comparing the two LTR sequences flanking each retrotransposon. As shown in Fig. 1b, the age distributions of *Copia* and *Gypsy* elements center on similar values (3.20 Million years (Myrs) vs. 3.37 Myrs, Mann-Whitney U test P >0.05). Both groups are younger than the unclassified retrotransposons that are on average 4.78 Myrs old (Mann-Whitney U test, both P<0.001).

Despite a similar median age, the *Gypsy* group has a much higher proportion of young copies compared to *Copia*. More than one third of *Gypsy* elements were less than two million years old (34.2%), whereas the *Copia* age distribution peaks around 3-4 Myrs with young copies (0-2 Myrs) accounting for only 23.9% (Kolmogorov-Smirnov test, p = 5.93×10^-^ 4; Fig. 1c). These results suggest that *Gypsy* elements have been active very recently in the *R. apiculata* lineage even though they did not reach the copy numbers exhibited by *Copia*.

The suppression of active TE copies is mainly achieved by *de novo* and maintenance DNA methylation. We therefore conducted bisulfite sequencing to examine genome-wide DNA methylation in *R. apiculata* leaves. LTR retrotransposons globally displayed a substantial increase in DNA methylation compared to their flanking regions (Fig. 2a). The increase in CG and CHG methylation was evenly distributed across whole TE bodies. In contrast, CHH methylation dramatically increased at TE boundaries that correspond to LTR regions (Fig. 2a). The difference was more than two-fold in the *Gypsy* group (19.9% in LTRs vs. 8.3% internally), with a more modest difference in *Copia* (13.9% in LTRs vs. 9.7% internally), and almost unnoticeable in the unclassified elements (2.9% in LTRs vs. 2.8% internally; Fig. 2a). Such differences in methylation could not be attributable to differential occurrence of CG, CHG and CHH sites in LTRs *versus* internal regions among LTR retrotransposon groups (Fig. S1). The low level of methylation in unclassified elements is consistent with previous observations (Bousios *et al.*, 2016) that older TEs are typically methylated less densely than younger ones.

**Fig. 2.**
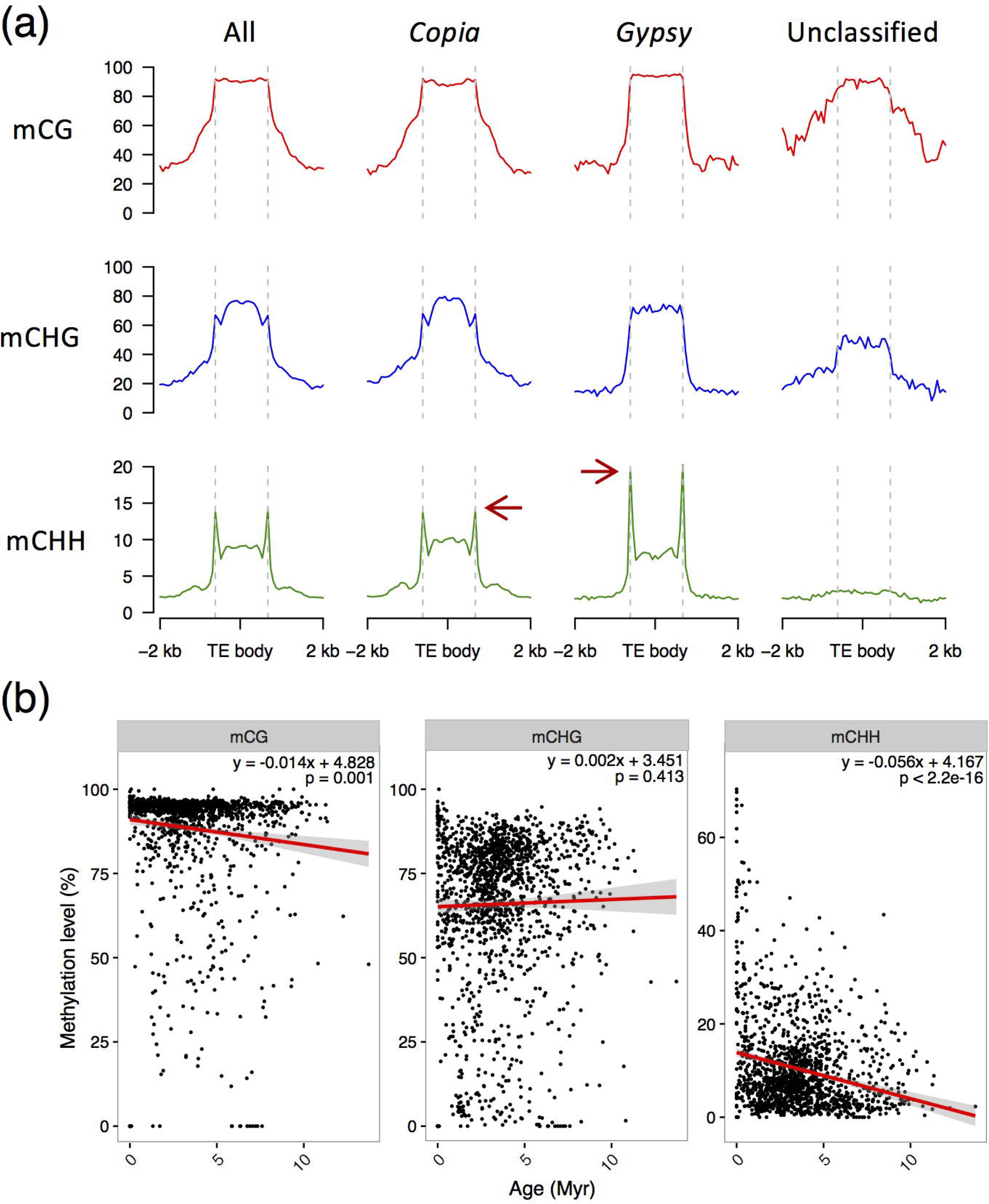
LTR retrotransposon methylation patterns in *R. apiculata*. (a) Methylation levels of LTR retrotransposons and their flanking regions. Each retrotransposon was divided into 20 equally-sized windows, in addition to two kilobases (kb) of the upstream and downstream flanking regions. The average methylation level for each window was calculated and plotted in red (CG), blue (CHG), and green (CHH). Dashed lines indicate TE body boundaries. Red arrows indicate the increase of methylation at TE ends that correspond to LTRs. (b) Correlations between CG, CHG, or CHH levels and LTR retrotransposon age. Red lines indicate regression curves and shadows delimit 95% confidence intervals.

CHH methylation is *de novo* DNA methylation that depends on the RNA-directed DNA methylation (RdDM) pathway (Zemach *et al.*, 2013; McCue *et al.*, 2015). It is therefore intriguing that this modification is concentrated in LTR regions that contain the promoters for retrotransposon transcription. This raises the possibility that newly-formed elements are being suppressed via CHH methylation in the *R. apiculata* genome. Indeed, we observe that young TEs are more heavily targeted by CHH methylation than older copies (Pearson’s correlation, *r* = - 0.23, P < 2.20×10^−16^; Fig. 2b). Although young *Gypsy* elements may retain an active transposition capability, the proliferation of these TEs may have been counteracted with strong CHH methylation in their LTRs.

### siRNA targeting associates with TE methylation

To understand the potential role of siRNAs in differential accumulation of TEs, we sequenced two replicate small RNA libraries from *R. apiculata* leaves. Short reads between 18 and 30 nucleotides (nt) in length were mapped to the reference genome of this species allowing no mismatches. After adapter trimming, quality control, and filtering of tRNAs, rRNAs, snoRNAs and known miRNAs, we obtained about 110,000 ± 10,000 distinct species of 21-nt siRNAs, 60,000 ± 3,000 of 22-nt, and 150,000 ± 14,000 of 24-nt siRNAs. We measured siRNA expression at LTR retrotransposons as read counts per base pair per element. Unless otherwise specified, if an siRNA matched multiple places in the genome, we assigned the total read count divided by the number of loci to each location and added these data points to the uniquely mapping siRNAs.

24-nt siRNAs are the most abundant species in *R. apiculata* whether they map to LTRs or TE bodies (Fig. 3a). Pooling siRNAs of all lengths, those mapping to LTR regions are significantly more abundant than those at internal TE regions (Mann-Whitney U test, all P values < 0.01, Fig. 3b). Targeting of *Gypsy* elements (median: 13.84 counts per nucleotide per element) is four to eight times more intense than association with *Copia* (3.11) or unclassified (1.61) elements (Fig. 3b). Furthermore, we found a robust positive correlation of siRNA levels and CHH methylation for all siRNA lengths (Partial correlation controlling for CG and CHG methylation, *r* = 0.19-0.30, all P < 1.48e-13; Fig. S2). Taken together, these results suggest strong RdDM activity associated with high CHH methylation accounts for the observed low abundance of *Gypsy* elements in *R. apiculata*. Given that *Gypsy* and *Copia* elements have similar ages on average (Fig. 1b), it is unlikely that *Gypsy* elements are too young to achieve a high copy number.

**Fig. 3.**
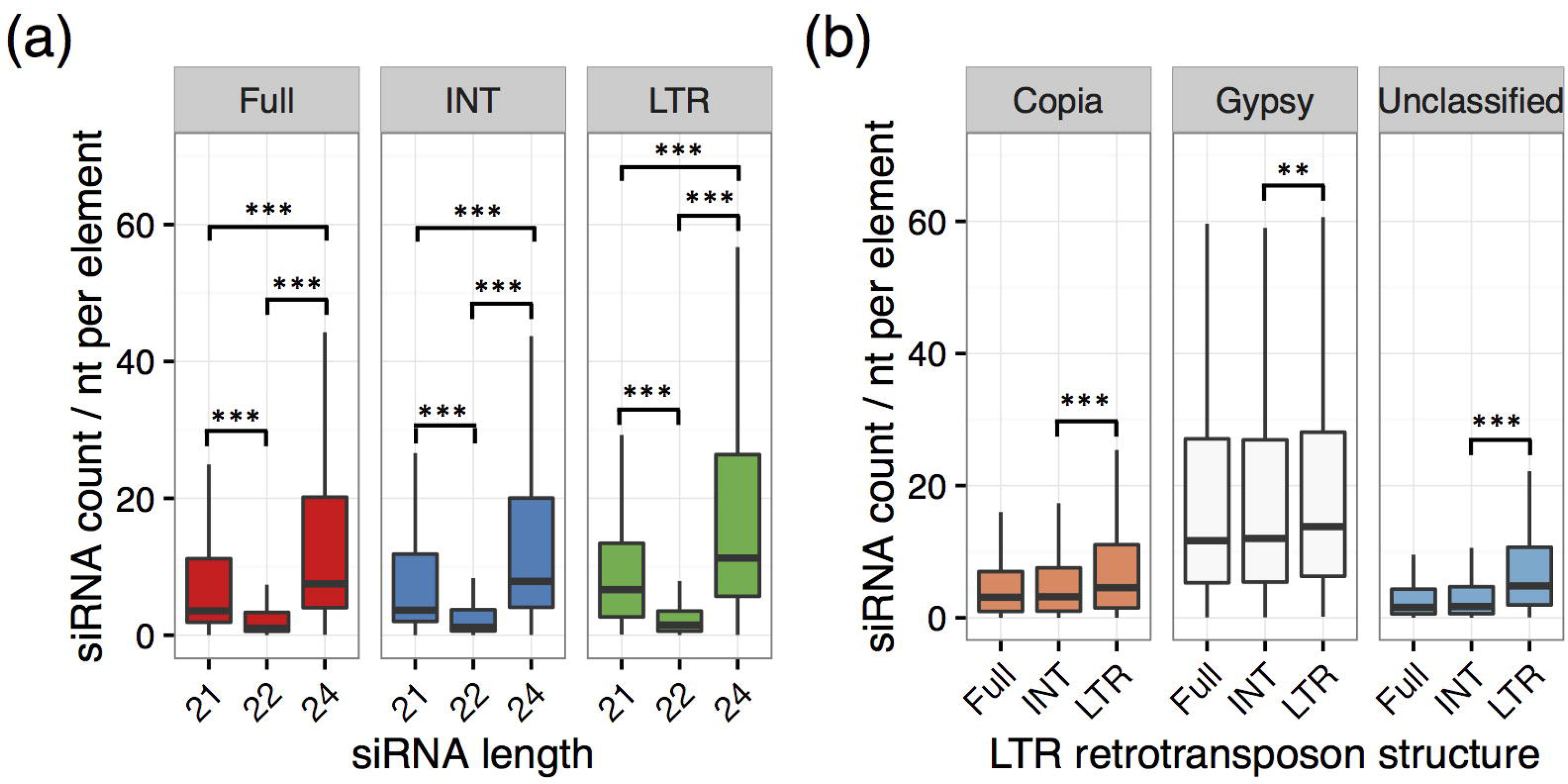
Patterns of siRNA targeting of *R. apiculata* LTR retrotransposons. (a) siRNA levels according to siRNA length and transposon region targeted. (b) siRNA levels by LTR retrotransposon group. A Mann-Whitney U test was used for comparison. P values are indicated as *: *P*<0.05, **: *P*<0.01, and ***: *P*<0.001.

### siRNA targeting of old *Gypsy* elements

Given that CHH methylation preferentially targets young elements, especially those from the *Gypsy* group, we wondered whether siRNA targeting shows the same age preference. To answer this question, we grouped TEs into age bins, separated by siRNA length and retrotransposon group. Fig. 4a shows that siRNA density at LTR retrotransposons drops for *Copia* and *Gypsy* elements that are more than one million years old (Fig. 4a). However, *Gypsy* elements of all ages harbor more siRNAs regardless of their length (Fig. 4a). When grouping siRNAs by the age of targeted TEs, the cumulative distribution for siRNAs at *Copia* elements is steeper than for those at *Gypsy* elements (Kolmogorov–Smirnov test, P < 2.20×10^−16^, Fig. 4b), suggesting a less uniform representation of ages among *Copia* elements targeted by siRNAs. Indeed, siRNAs mapped to *Gypsy* copies older than four million years account for 35.1% of the total counts, while the fraction is only 25.1% of the *Copia* elements. The high siRNA density at old *Gypsy* elements is unusual because old elements (as is evident among the unclassified TEs) typically lose siRNA targeting due to mutation accumulation (Fig. 4a). Our results thus suggest old *Gypsy* elements tend to be reserved or reactivated to produce siRNAs.

**Fig. 4.**
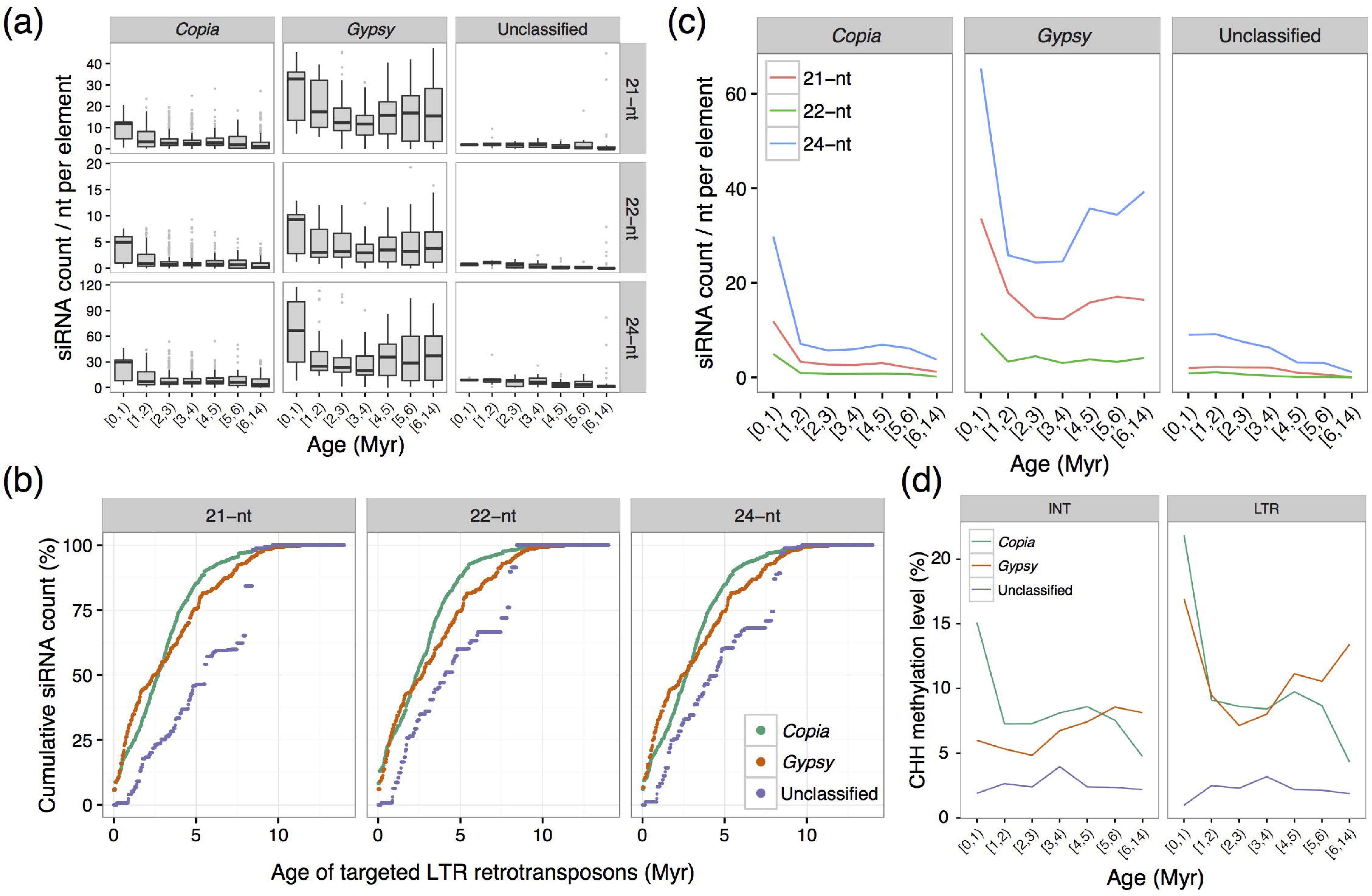
siRNA targeting and TE age. (a) siRNA levels at LTR retrotransposons by age bin. (b) Cumulative plot of siRNA levels at LTR retrotransposons according to age. (c) Median level of siRNA expression at LTR retrotransposons from different age bins. (d) Median level of CHH methylation at LTR retrotransposons from different age bins.

It is surprising that siRNA density at old *Gypsy* elements actually increased when we looked at medians within each age group introduced in panel A of Fig. 4. While all siRNA activity sharply drops for *Copia* elements older than two million years, there is a clear increase in 21-nt and 24-nt siRNAs at *Gypsy* elements older than four million years (Fig. 4c). The 21/22-nt siRNAs mediate post-transcriptional silencing (PTGS) and act as the first layer of defense against newly-emerging TE copies (Mari-Ordonez *et al.*, 2013). The 24-nt siRNA-mediated transcriptional silencing (TGS) then takes over as TEs accumulate (Mari-Ordonez *et al.*, 2013). The unusual increase of siRNAs suggests a considerable number of *Gypsy* elements have undergone a kind of silencing rejuvenation.

To test this hypothesis, we compared the relationship between CHH methylation and TE age among different LTR retrotransposon groups. As expected, the level of CHH methylation increased at old *Gypsy* elements, especially in the LTR regions (Fig. 4d). In contrast, old *Copia* elements exhibited a much lower level of CHH methylation which decreased with TE age (Fig. 4d). Experimental data have demonstrated that silencing of active TEs occurs quickly, within a few host generations (Teixeira et al., 2009; Mari-Ordonez et al. 2013; Fultz and Slotkin 2017). It is thus very likely that old *Gypsy* elements have been silenced at least once and then reactivated more recently.

To test whether the unusual age spectrum of siRNA targeting involves particular TE family, we separated TEs into young (0-4 Myrs) and old (>4 Myrs) groups and compared siRNA densities between groups within each family that contained more than three copies (Fig. S3a). Only two out of the 18 families in our survey showed significantly higher siRNA expression at old TE copies: the *Gypsy* family RapLTR32 and the *Copia* family RapLTR11 (Fig. 5a). There were 30 intact elements in RapLTR32 and 74 in RapLTR11. The mean number of siRNAs mapping to RapLTR32 is three times greater at the old (94.77) elements than at young insertions (33.09, Mann-Whitney U test P = 3.21×10^−6^). This is in striking contrast to the genome-wide pattern (9.94 siRNA copies at old vs. 13.00 at young, Mann-Whitney U test P = 2.17×10^−10^). Although the overall siRNA expression at RapLTR11 is low (4.92 siRNA copies at old vs. 2.24 at young), the pattern is the same as for RapLTR32 (Mann-Whitney U test, P = 5.33×10^−5^).

**Fig. 5.**
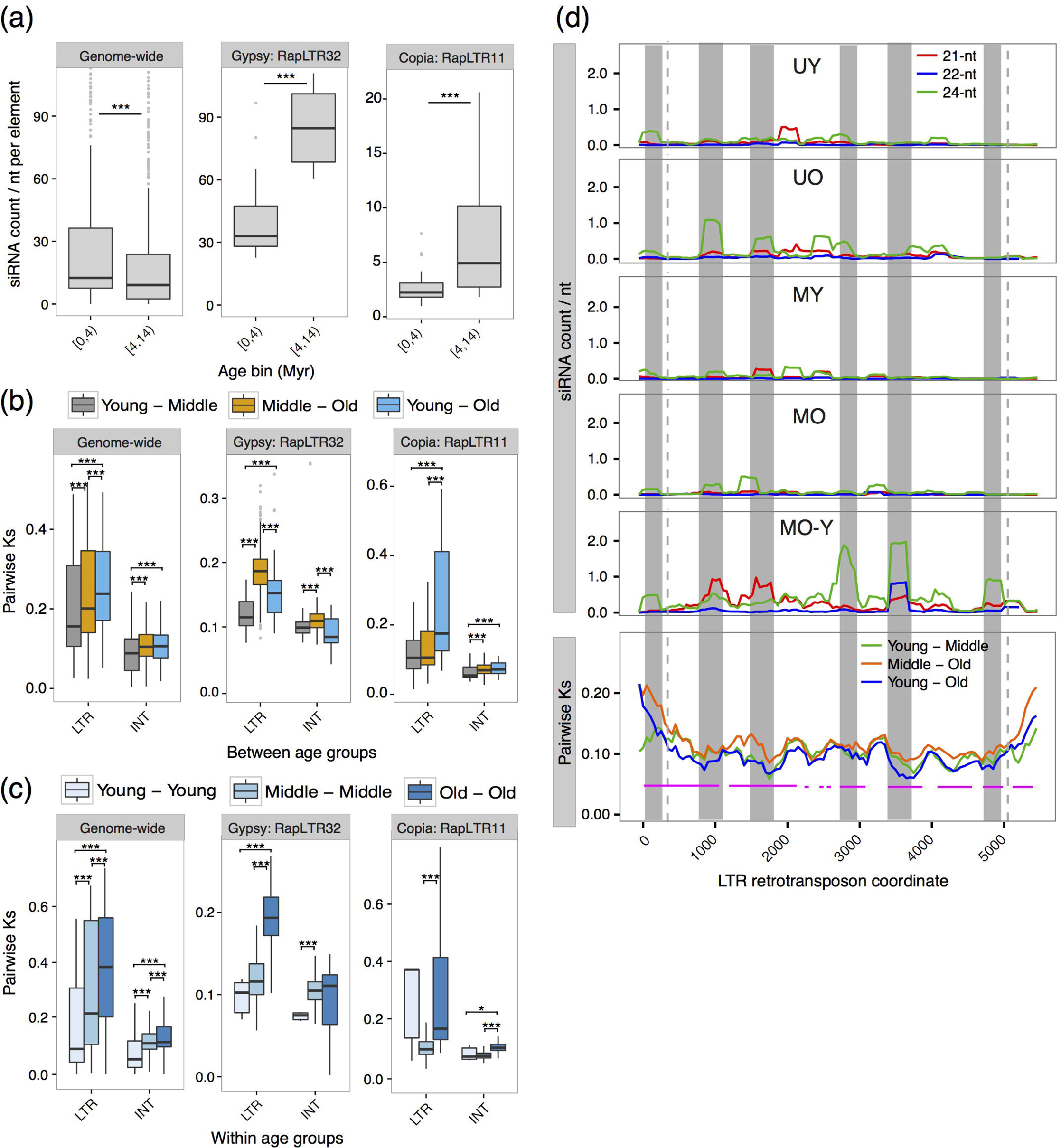
Comparative analyses of RapLTR32, RapLTR11, and all *R. apiculata* LTR retrotransposons. (a) siRNA expression at the young (0-4Myrs) and old (>4 Myrs) elements. (b) Pairwise Ks between groups of young (0-2Myrs), middle-aged (2-4Myrs), and old (>4 Myrs) elements. (c) Ks within young, middle-aged, and old elements. A Mann-Whitney U test was used for comparison. *P* values are indicated as *: *P*<0.05, **: *P*<0.01, and ***: *P*<0.001. (d) Sliding window analysis of siRNA level and sequence divergence in RapLTR32. Window size is 250 bp and step width is 50 bp. The upper panel shows levels of 21-nt (red), 22-nt (blue), and 24-nt (green) siRNAs. The bottom panel shows average Ks between young and middle-aged elements (green), middle-aged and old elements (orange), and young and old elements (blue). Pink lines indicate regions where Ks between young and old elements is significantly smaller than that between middle-aged and old elements (two-tailed *t* test with Benjamini–Hochberg correction, FDR< 0.05). Dash lines indicate the boundary between 5’ or 3’ LTR and the internal region. Shades indicate the overlaps between siRNA hot spots and low-divergence regions. UY, uniquely mapped to young elements; UO, uniquely mapped to old elements; MY, multiply mapped to young elements; MO, multiply mapped to old elements; MO-Y, cross-mapped between young and old elements.

### siRNA targeting cross-talk between young and old TEs

We wondered why old *Gypsy* elements harbor a high siRNA density. Previous studies suggested that the reactivation of silenced TEs can trigger *trans*-silencing of active relatives (Lisch, 2009; Bousios et al., 2016). It is possible that some old *Gypsy* elements became zombies and were retained as immunity memory in the *R. apiculata* genome. To test this hypothesis, we traced which TE age bin the siRNAs at old copies come from. As mutations accumulated with TE age, old copies tended to diverge and therefore to harbor more uniquely mapping siRNAs (U-siRNAs), whereas young copies are prone to harbor multiply mapping siRNAs (M-siRNAs) due to high sequence similarity among insertions. Indeed, TE age negatively correlates with M-siRNA density (Fig. S4a,c) but positively correlates with U-siRNA density (Fig. S4b,d). While siRNAs specific to old TEs (both U- and M-siRNAs) did account for a larger fraction of *Gypsy* elements (23%) than *Copia* elements (12%), around half of siRNAs cross-mapped between young and old TEs. These cross-mapping molecules contribute substantially to the siRNA density at old elements (Table 2).

**Table 2.** Multiply mapping siRNAs at RapLTR32, RapLTR11, and LTR retrotransposons genome-wide. TE copies were classified as young (0-4Myrs) or old (>4 Myrs). Numbers shown are the siRNA counts. The counts of multiply mapping siRNAs were weighed by # hits at young or old TE/ # total hits.

|  | Uniquely mapping |  | Multiply mapping |  |  |  | Total |
| --- | --- | --- | --- | --- | --- | --- | --- |
|  | Young | Old | Only young | Only old | Young and old |  |  |
|  |  |  |  |  | Young | Old |  |
| *Copia* | 13,177 (7%) | 13,129 (7%) | 47,374 (24%) | 10,496 (5%) | 98,461(49%) | 18,549(9%) | 201,186 |
| *Gypsy* | 14,128 (10%) | 17,365 (12%) | 30,318 (21%) | 16,437 (11%) | 56,805(38%) | 12,798(9%) | 147,850 |
| Unclassified | 633 (7%) | 4,500 (49%) | 765 (8%) | 1,883 (20%) | 1,143(12%) | 353(4%) | 9,277 |
| RapLTR32 | 717 (5%) | 2,774 (18%) | 2,092 (14%) | 1,807 (12%) | 4,501(29%) | 3,554(23%) | 15,445 |
| RapLTR11 | 109 (2%) | 415 (6%) | 1,295 (18%) | 2,812 (40%) | 1,851(26%) | 587(8%) | 7,069 |

If selection favors cross silencing of new elements by old elements, we expect low sequence divergence between young and old *Gypsy* copies, and substantial expression of old *Gypsy* elements. We calculated pairwise *Ks* between groups of young (0-2 Myrs), middle-aged (2-4 Myrs) and old (>4 Myrs) elements from different LTR retrotransposon groups (Fig. 6a). Ks between young and old elements was indeed smaller than Ks between middle-aged and old elements (Mann-Whitney U test, P<0.001), but only in the internal regions. To measure TE expression, we conducted an RNA-seq analysis using *R. apiculata* leaves. While most TEs were barely expressed, expression level increased with *Gypsy* element age (Pearson’s correlation, *r*= 0.49, P< 0.001, Fig. 6b). Notably, old *Gypsy* elements exhibited measurable expression, as expected.

**Fig. 6.**
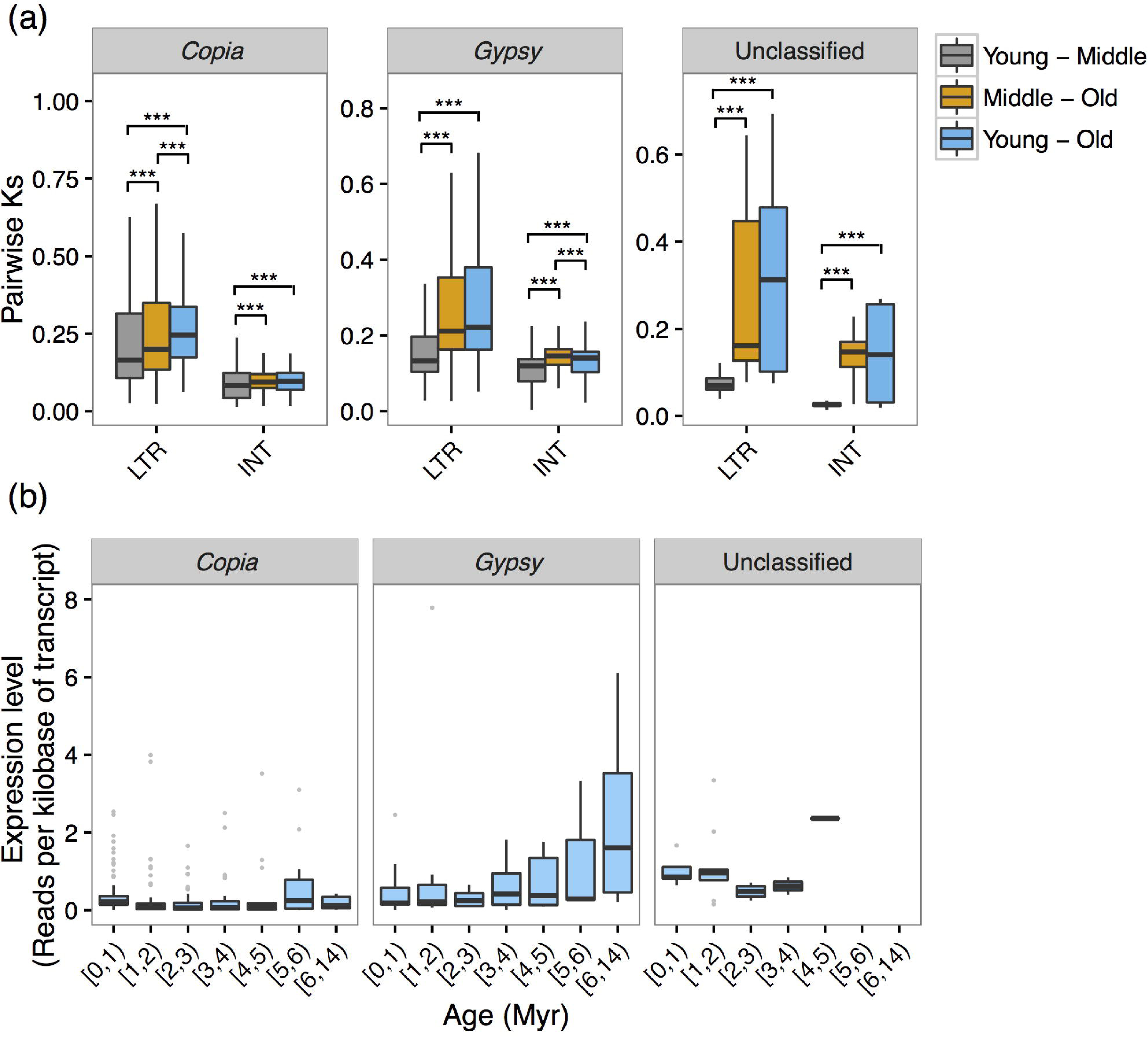
Sequence divergence and expression level of LTR retrotransposons from young (0-2 Myrs), middle-aged (2-4 Myrs) and old (>4 Myrs) age groups. (a) Pairwise *Ks* between groups of young, middle-aged and old elements. A Mann-Whitney U test was used for comparison, and *P* values are indicated as *, *P*<0.05; **, *P*<0.01; and ***, *P*<0.001. (b) Expression level of LTR retrotransposons from different age bins. Expression level of each LTR retrotransposon copy was calculated as Reads Per Kilobase per Million mapped reads (RPKM).

We then focused on *Gypsy* RapLTR32 and *Copia* RapLTR11 families, exhibiting surprisingly prevalent high siRNA density at old copies, and checked the impact of siRNA targeting cross-talk between young and old elements. RapLTR32 show a much higher proportion (23%) of the young-and-old cross-mapping siRNAs at old copies than *Gypsy* elements overall (9%), whereas the proportions in RapLTR11 are comparable to *Copia* group (8% vs. 9%, Table 2). Correspondingly, RapLTR32 shows a low divergence between young and old copies (Fig. 5b, middle panel), especially compared to within-age-group differentiation (Fig. 5c, middle panel). In contrast, the old/young divergence is high among RapLTR11 elements (Fig. 5b,c, right panels). As a control, between- and within-age-group differentiation increases with TE age genome-wide (Fig. 5b,c, left panels) and for most TE families (Fig. S3b,c), particularly at LTR sites. A sliding window analysis of RapLTR32 revealed hot spots for producing siRNAs that allow silencing of young elements by old TEs. These regions exhibited particularly low divergence between these age groups (Fig. 5d).

Taken together, our results indicate that old *Gypsy* elements were selectively retained in the *R. apiculata* genome, and reactivation of these zombies led to production of siRNAs triggering *de novo* silencing of subsequent waves of new elements.

### TE effect on host gene expression and its evolutionary consequences

Transcriptional silencing of TEs can potentially affect the expression of neighboring genes by establishing repressive epigenetic markers (Hollister & Gaut, 2009). We expected that *Gypsy* elements would have stronger deleterious effect on the expression of nearby genes, which in turn would result in intense selection against their accumulation.

We first calculated distances between TE insertions and closest annotated genes in the *R. apiculata* genome. *Gypsy* elements are indeed on average closer to genes than *Copia* elements (Fig. 7a). This relationship holds for both the young (0-4 Myrs) and old (>4 Myrs) copies, although it is statistically significant only between the latter (Mann-Whitney U test P < 0.01). Interestingly, old insertions of both groups are typically slightly closer to genes than new elements (Mann-Whitney U test, P < 0.05; Fig. 7a), although a finer age stratification reveals that the median distance to genes peaks at an intermediate age (Fig. S5a).

**Fig. 7.**
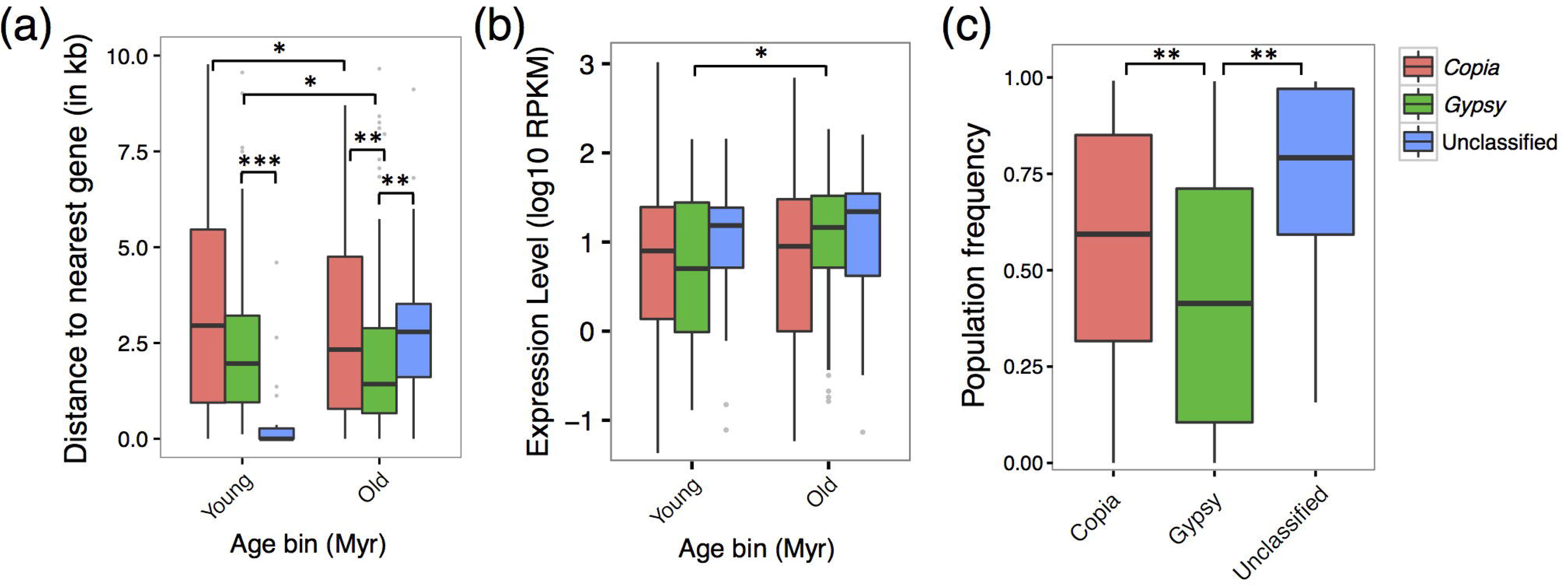
TE effects on host gene expression and their evolutionary consequences. (a) Distance between the young (0-4Myrs) and old (>4 Myrs) elements from three LTR retrotransposon groups and nearest genes. A distance of zero indicates the TE is located within the intron or UTR of a gene. (b) Expression of genes nearest to the young (0-4Myrs) and old (>4 Myrs) elements. Gene expression was calculated as Reads per Kilobase per Million mapped reads (RPKM). (c) Population frequency of TE insertions of *Copia*, *Gypsy* and unclassified TEs. A Mann-Whitney U test was used for comparison. *P* values are indicated as *: *P*<0.05, **: *P*<0.01, and ***: *P*<0.001.

Using RNA-seq data, we measured the potential effect of transposon insertions on expression of nearby genes. Median expression levels of genes located close to young *Gypsy* elements is indeed slightly lower than that of genes neighboring *Copia* elements (Fig. 7b). Furthermore, loci next to young *Gypsy* insertions are typically under-expressed compared to genes near old elements of that group (Mann Whitney U test P < 0.05, Fig. 7b). In contrast, genes near old and young *Copia* elements are at comparable expression levels (Mann Whitney U test P > 0.05, Fig. 7b). These results should be interpreted with caution because TE insertions may sometimes provide promoters or enhancers that induce downstream gene expression (Feschotte, 2008).

To estimate selection intensity, we conducted high-throughput sequencing of a pool of ten *R. apiculata* individuals and estimated population frequencies of TE insertions. The expectation is that deleterious elements will fail to reach high frequency in the population. After filtering to eliminate low-confidence calls (see Materials and Methods), we identified 1,439 out of the 1,446 insertions considered in this study so far. Out of these, 238 elements were polymorphic in our population of ten individuals. These include 148 *Copia* insertions, 77 *Gypsy*, and 11 unclassified. Consistent with the possible deleterious effects of *Gypsy* gene proximity, we see a significantly lower allele frequency among TEs of this group compared to *Copia* and unclassified elements (Mann-Whitney U test, both P < 0.01, Fig. 7c). We further confirmed our allele frequency estimates by screening a panel of 32 *R. apiculata* individuals for presence/absence of 23 randomly chosen TE insertions using a PCR-based assay (Fig. S6). The population frequencies estimated using the two methods were highly concordant (Pearson’s correlation among allele frequencies, r = 0.79, P = 6.27×10^−6^). Taken together, our results suggest *Gypsy* insertions are highly deleterious and under strong purifying selection.

## Discussion

The well-known genome shock hypothesis suggests that stress-induced reactivation of TEs can lead to genome rearrangements that in turn may facilitate genetic adaptation (McClintock, 1984). Convergent genome size reduction observed in mangroves apparently contradicts this hypothesis. It is thus puzzling how mangroves reconcile the conflict between stress-induced TE activation and selection for genome size reduction. This reduction is thought to be due to a dampening of LTR retrotransposon transposition rates (Lyu *et al.*, 2018). Indeed, we found higher Ts:Tv ratio in *R. apiculata* than in non-mangroves (Table 1), suggesting TE activity in this species is suppressed by strong DNA methylation. However, the *Gypsy* group of LTR retrotransposons remained active very recently within the *R. apiculata* genome. Not only are young *Gypsy* elements preferentially methylated at CHH sites (Fig. 2), but the old TEs from this group are heavily targeted by 21- and 24-nt small RNAs (Fig. 4). Therefore, our results suggest a continuous battle between TEs and the mangrove host obscured by the observed low TE load.

It is unclear why these particular TE families have undergone apparent reactivation. One possibility is that some of these TEs have been introduced via horizontal transfer. Although such events have been previously reported in mangroves *Excoecaria agallocha* and *Kandelia candel (Liu et al., 2016; Huang et al., 2017)*, we found the *Gypsy* and *Copia* group are at similar age on average, which does not seem to support this possibility. Another plausible explanation is a reactivation of existing TEs due to genomic shock, e.g. hybridization, polyploidy, and stress (Madlung & Comai, 2004; Madlung *et al.*, 2005; Kenan-Eichler *et al.*, 2011). Several TEs, such as *Tnt1* in tobacco (Grandbastien *et al.*, 1997), *ONSEN* in Arabidopsis (Cavrak *et al.*, 2014; Gaubert *et al.*, 2017), and *Tos17* in rice (Hirochika *et al.*, 1996) are activated by changes in the environment. The increase in genetic diversity due to stress-induced transposon activation may increase the capacity of species to evolve stress resistance traits (Kidwell & Lisch, 1997; Horvath & Slotte, 2017). This scenario is particularly intriguing in mangroves because these species harbor extremely low standing genetic variation (Guo *et al.*, 2017) but live in stressful habitats that require constant adaptation. Additional transposition events can thus increase the evolutionary potential of these species. The TE composition and methylation patterns in *R. apiculata* we observed may be a result of a balance between high TE activity and stringent epigenetic regulation that keeps most deleterious elements at bay. Indeed, we observe that old *Gypsy* elements, the ones that have been maintained in the face of more than four million years of selection and drift, are expressed even more (Fig. 6b) and exhibit considerable CHH methylation (Fig. 4d). In contrast, unclassified elements, which are usually old, are poorly targeted by CHH methylation and siRNAs, and show much higher population frequency than *Copia* and *Gypsy* elements, suggesting they are better tolerated by the host likely due to loss of retrotransposition capacity.

A surprisingly large fraction of siRNAs that map to several loci in the *R. apiculata* genome target both old and new copies. This suggests that old elements, even if they only produce partial transcripts, may act as reservoirs of silencing information that can be deployed against newly emerging insertions (Lisch, 2009). This scenario seems particularly likely in the *Gypsy* RapLTR32 family, where divergence between young and old elements is unusually low (Fig. 5). In RapLTR312, about half of the cross-mapping siRNAs between young and old copies reside at the old elements (Table 2). This result indicates the reactivation of so-called “zombie” copies can rapidly trigger silencing *in trans* of young elements from the same family. Similar hypotheses have been proposed for the tobacco *Tnt1* element (Perez-Hormaeche *et al.*, 2008), maize Sirevirus elements (Mari-Ordonez *et al.*, 2013), and heterochromatic clusters of TEs in animals (Aravin *et al.*, 2007; Brennecke *et al.*, 2007).

Silencing of LTR retrotransposons has deleterious effects on nearby gene expression. Heavily methylated *Arabidopsis* TEs are more likely to be quickly removed from the genome if they reside near genes (Hollister *et al.*, 2011). We do observe that *Gypsy* elements are more likely to reside close to genes than *Copia* TEs. However, they are at relatively low frequencies in the population we surveyed, suggesting a stronger deleterious effect. Perhaps *Gypsy* elements are more prone to land next to genes, and selection has not had time to purge all of them. We do observe that *Gypsy* elements suffer higher rates of internal deletion, as measured by comparing LTR and internal region lengths, than do *Copia* retrotransposons (Table S2). This suggests an increased selection pressure to purge *Gypsy* elements from the *R. apiculata* genome by recombination. Overall, we find that high origination rates and quick removal lead to a constant evolutionary flux of *Gypsy* elements in this mangrove species.

Our results also shed light on the factors that contribute to the process of epigenetic TE silencing. Palindromic structures in the maize Sirevirus LTRs can form hairpins that are processed into “primary” siRNAs and aid in silencing of active elements (Bousios *et al.*, 2016). Although we searched LTRs of *R. apiculata* transposons for secondary RNA structures, we did not find any TE families prone to form hairpins. The patterns of CHH methylation and siRNA targeting we see in the *R. apiculata* genome do suggest that LTRs may be the hot spot target for siRNA-mediated RdDM in this species (Fig. 2a and Fig. 3). However, experimental reconstruction of *de novo* silencing of active *EVD* copies in *Arabidopsis* revealed a PTGS-to-TGS shift, which was initiated by 21- to 22-nt 3’ *gag*-derived siRNAs and spread to 5’ LTR *de novo* methylation mediated by 24-nt siRNAs (Mari-Ordonez *et al.*, 2013). As such a transition occurs very quickly within several generations (Mari-Ordonez *et al.*, 2013), the LTR-derived 24-nt siRNAs in *R. apiculata* may represent the final stage rather than the initiation of the epigenetic control of active TE elements.

Taken together, our results indicate that the reactivation of TEs, possibly by stress, may cause rapid suppression of active elements in *R. apiculata*, forming a feedback loop that eventually leads to stable epigenetic control of TEs. By tackling the most dangerous elements, the constant arms race between TEs and the host continuously provides new genetic material that may facilitate stress adaptation in mangroves.

## Acknowledgements

We thank Anthony Greenberg for critical reading and language editing of the manuscript. We thank Shaohua Xu for help with the *R. apiculata* genome data. This study was funded by the National Key Research and Development Program of China (2017YFC0506100-1), the National Science Foundation of China (31770246), the Science and Technology Program of Guangzhou (201707020035), the Fundamental Research Funds for the Central Universities (16lgjc75), the program of Guangdong Key Laboratory of Plant Resources (PlantKF05), and the Chang Hungta Science Foundation of Sun Yat-sen University.

## Author contributions

T.T. and Y.W. planned and designed the research; Y.W. and W. L. conducted the experiments; Y. W. and T. T. analyzed the data; Y.W. and T.T. wrote the manuscript. All authors have read and approved the manuscript. The authors declare that the research was conducted in the absence of any commercial or financial relationships that could be construed as a potential conflict of interest.

## Supporting Information

Additional supporting information may be found in the online version of this article.

**Fig. S1.** Occurence of CG, CHG, and CHH sites in LTRs and internal regions of different LTR retrotransposon groups.

**Fig. S2.** Patterns of siRNA targeting and TE methylation levels of LTR retrotransposons in *R. apiculata*.

**Fig. S3.** Comparative analyses of all the chosen LTR retrotransposons in *R. apiculata*.

**Fig. S4.** Correlations between multiply or uniquely mapping 21-/22-nt or 24-nt siRNA targeting intensity and TE age.

**Fig. S5.** Correlation of TE age with distance to and expression of their nearest genes.

**Fig. S6.** PCR validation of the presence/absence status of LTR retrotransposon insertions.

**Table S1** List of all intact LTR retrotransposons in *R. apiculata.*

**Table S2** Copy number ratios of LTRs to internal regions in different groups of LTR retrotransposons in *R. apiculata*.

**Table S3** Primer pair combinations used to detect the presence and absence of LTR retrotransposon insertions in *R. apiculata*.

